# Repliscan: a tool for classifying replication timing regions

**DOI:** 10.1101/094177

**Authors:** Gregory J. Zynda, Jawon Song, Lorenzo Concia, Emily E. Wear, Linda Hanley-Bowdoin, William F. Thompson, Matthew W. Vaughn

## Abstract

**Background:** Replication timing experiments that use label incorporation and high throughput sequencing produce peaked data similar to ChIP-Seq experiments. However, the differences in experimental design, coverage density, and possible results make traditional ChIP-Seq analysis methods inappropriate for use with replication timing.

**Results:** To accurately detect and classify regions of replication across the genome, we present Repliscan. Repliscan robustly normalizes, automatically removes outlying and uninformative data points, and classifies Repli-seq signals into discrete combinations of replication signatures. The quality control steps and self-fitting methods make Repliscan generally applicable and more robust than previous methods that classify regions based on thresholds.

**Conclusions:** Repliscan is simple and effective to use on organisms with different genome sizes. Even with analysis window sizes as small as 1 kilobase, reliable profiles can be generated with as little as 2.4x coverage.

## Background

The most essential property of the cell is its ability to accurately duplicate its DNA and divide to produce two daughter cells [1]. The cell’s replication cycle starts with G1 phase, in which molecules essential for cell division are produced, then proceeds to replicating DNA in S phase. After all DNA in the genome is duplicated, the cell continues to grow in G2 phase until it divides into two daughter cells at the end of Mitosis, or M phase, at which point it is ready to start the cell cycle again (Figure 1).

To ensure accuracy and efficiency, S phase is complex and highly regulated. Instead of duplicating in a single zipping motion, reminiscent of transcription, DNA is synthesized in regions at distinct times in eukaryotes, initiating at multiple origins of replication [2]. This synthesis process takes place in a live cell, so replication mechanisms need to be coordinated with active transcription, chromatin configuration, and three-dimensional structure [3]. For example, early replication correlates with chromatin accessibility [4].

To better understand the coordinated program of DNA replication, two types of protocols have been developed to examine genome-wide replication profiles based on DNA sequencing data. One based on the time of replication, TimEx [5, 6], and the other based on incorporation of a labeled precursor into newly replicated DNA, Repli-seq [7, 8, 9, 10, 11, 12]. Time of replication (TimEx) measures DNA coverage at sequential times in S-phase. The normalized early S-phase signal should be mostly 1x coverage, additively transitioning to 2x coverage in late S-phase. In contrast to this method, Repli-seq works by only sequencing newly replicated DNA. Theoretically, in a single cell, this means once a region is replicated, it should not appear in samples taken at later times, except in the case of allelic timing differences. Both methods have been shown to yield similar results [13, 14] for when and where genomic regions replicate, but each requires a distinct type of analysis. The methods described in this paper focuses on data produced by label incorporation (Repli-seq).

**Figure 1.**
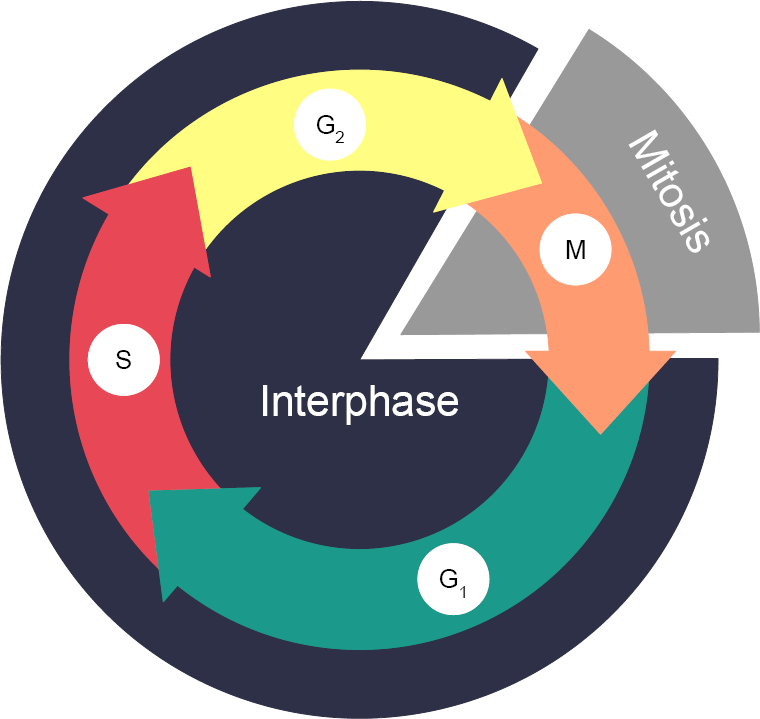
Overview of the cell cycle. Cell division takes place in two stages: interphase and mitosis. Interphase is when a cell copies its genome in preparation to physically divide during mitosis. Interphase starts with cell growth and preparation for DNA synthesis in Gap (G1). After G1, DNA is replicated in regions during the Synthesis (S) phase. The cell then transitions into a second growth phase – Gap 2 (G2). When the cell has finished growing, the cell divides into two daughter cells in Mitosis (M).

### Data Description

In continuation to our analysis of *A. thaliana* chromosome 4 in 2010 [15], we updated our laboratory protocol to be more stringent as described in Hansen *et al.* 2010[12], Bass *et al.* 2014[16], Bass *et al.* 2015[17], and Wear *et al.* 2016[18]. We increased the sensitivity of the labelling process by using 5-Ethynyl-2’-deoxyuridine (EdU), which does not require harsh denaturation of DNA, unlike 5-Bromo-2’-deoxyuridine (BrdU) used in previous work. A flow cytometer is then used to separate labeled from unlabeled nuclei, and to resolve labeled nuclei into different stages of S phase based on their DNA content. Next, DNA is extracted from sorted nuclei. The newly replicated DNA is immunoprecipitated and then sequenced using an Illumina sequencer. Previous protocols used microarrays for labeled DNA detection, which provided signal on probes at fixed intervals across a genome. Directly sequencing the immunoprecipitated DNA allows for a continuous display of replication activity across the genome.

Following the Repli-seq protocol, we created an exemplar *A. thaliana* dataset for development, with nuclei from: G1 (non-replicating control) and early, middle, and late S phase. While the amplification, fragmentation, and sequencing of next generation sequencing (NGS) libraries should be unbiased and random, physical factors affect the sequenceability of each region. To correct for these effects, we use the raw non-replicating DNA from the G1 control to normalize any sequenceability trends.

### Introducing Repliscan

In addition to our updated laboratory protocol for generically measuring DNA replication, we needed to improve the sensitivity and robustness of our analytical method. In previous work, log-ratios and aggressive smoothing were used to classify genomic regions by their time of replication. While this yielded results with high true positive rates, we found that this approach over-smoothed our deep coverage, next generation sequencing data. We created the Repliscan method to analyze generic, DNA sequence-based replication timing data without user-specified thresholds. Accepting any number of S-phase timepoints as input, Repliscan removes uninformative or outlying data, smooths replication peaks, and classifies regions of the genome by replication time.

## Methods

The analysis of the replication time data starts like any other DNA sequencing analysis, with quality control, mapping, and alignment filtering. Quality control consisted of removing contaminating 3’ universal sequencing adapters from the paired reads, and trimming the 5’ ends with quality scores below 20 with the program Trim Galore![19] version 0.3.7, which is designed to maintain read pairs. While it is obvious that low-quality regions need to be removed or masked because those base calls are untrustworthy, any contaminating sequences from adapters hinder the alignment process even more because they are always high-quality and may comprise a large part of the read. Therefore, reads in the output from Trim Galore! shorter than 40 base pairs were discarded, and resulting singletons (unpaired reads) were not included for alignment.

We then used BWA-MEM [20] version 0.7.12 with default parameters to align the quality-filtered reads to the TAIR10 *A. thaliana* reference genome [21]. After alignment, we filtered out any reads with multiple alignments using samtools[22] version 1.3. Removing these non-uniquely aligning reads is essential because they come from repetitive elements or other duplications in the genome that could replicate at different times, thereby confounding region classification into discrete replication times. After our stringent alignment requirements, fewer than 0.5% of our reads were identified as duplicates by samtools. We decided that removing the duplicates from our data was unnecessary due to the depth of our sequencing and localized nature of replication peaks. We also performed a correlation analysis of our samples and replicates, confirming their high level of similarity.

### Windowing

The DNA sequencing workflow leaves us with raw replication signals across a genome, which we must classify into distinct genomic regions and assign replication times. Our methods for this process build on methods from Lee *et al.* [15] and are illustrated in Figure 2.

**Figure 2.**
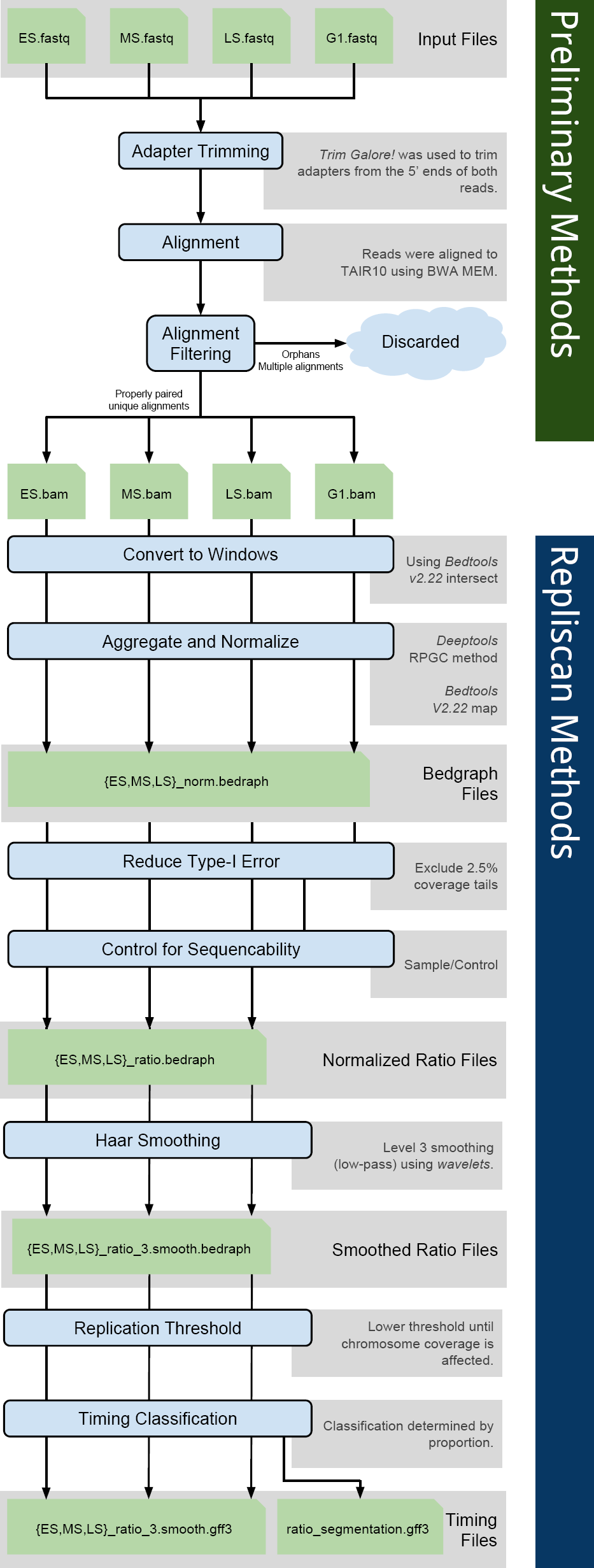
Repliscan workflow. Diagram of the preliminary alignment and quality control methods at the top, and the Repliscan methods at the bottom.

At first glance, Repli-seq data appears similar to dense ChIP-seq data [23], when viewed in a genome browser (Figure 3). However, instead of highlighting a limited number of coverage peaks as sites of molecular interactions, replication timing data consists of coverage across the entire genome accented with extremely wide peaks corresponding to regions of replication initiation and subsequent spreading. This background coverage with subtle, broad increases in depth makes deep coverage essential to reduce sampling error when detecting statistically-relevant differences. Even though the cost of sequencing has plummeted since 2007, deep-coverage DNA sequencing is still expensive for higher eukaryotes.

Lee *et al.* defined putative replicons in *A. thaliana* and calculated the median length to be 107 kilobases [15]. To achieve greater signal depth in each replication timing sample, we transformed each BAM alignment file into 1 kilobase coverage windows using bedtools [24]. While this transformation slightly reduces the resolution of our analysis, Figure 3 shows that the proportion of sampling error to measured signal is greatly reduced with the increased coverage. The windows also put all changes in coverage on the same coordinate system, simplifying comparisons between samples and experiments.

**Figure 3.**
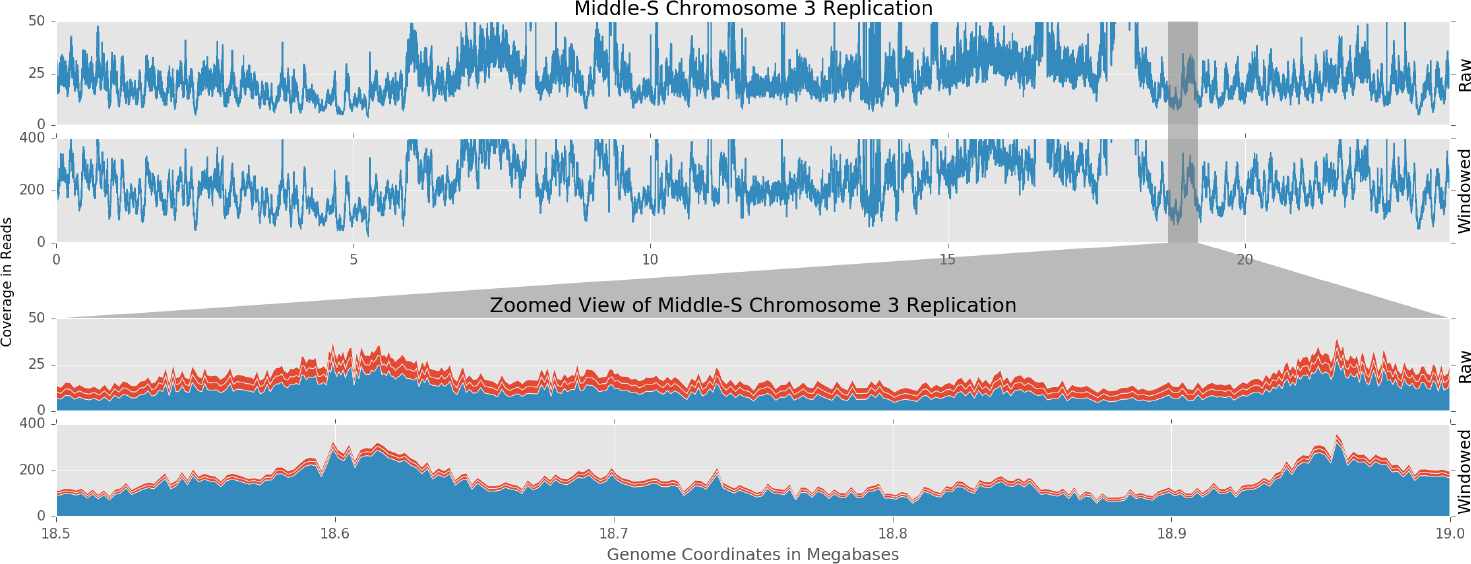
Replication signal and sampling uncertainty. The top two graphs show raw and windowed replication signal across *A. thaliana* chromosome 3. The bottom two graphs show raw and windowed replications signals at 18.5-19.0 megabases from the top view as represented by the gray selection area. The red bars represent sampling uncertainty (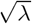 for Poisson distributions).

We chose 1 kilobase windows because they not only reduce sampling error, but are also two orders of magnitude smaller than the expected *A. thaliana* replicons. Repliscan does not summarize information with sliding windows, so choosing a window size that is an order of magnitude smaller than the expected replicon size is important to approximately align to the actual replication borders. Our analysis will theoretically allow the detection of regions of replication as small as 1 kilobase; however such regions are unlikely to exist in cells subjected to realistic labeling protocols. Therefore, in the final timing classification, Repliscan will merge neighboring regions with similar properties into larger segments. The 1 kilobase resolution then helps to highlight transitions between such segments. In some circumstances, such as working with low coverage data, it may be advantageous to use a larger window size. However, to achieve the best results when adapting Repliscan to other species, we suggest the expected replicon size be factored into calculations that establish window size and sequencing depth.

### Replicate Aggregation and Normalization

To further decrease sampling effects, and achieve consistent results between experiments, we used multiple biological replicates and adopted aggregation methods to either increase coverage or summarize replication signals using functions provided by “bedtools map” [24]. For experiments with low coverage, we pooled timing *t* = 1..*T* replicates *r* = 1..*R* together by summing coverage signal *k* across each window *i* = 1..*N*.

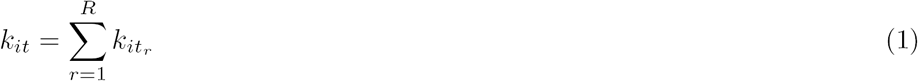

When coverage was sufficient, we used the signal mean (or the more robust signal median) to clean up aberrant coverage. For these methods, replicates were first normalized for sequencing depth using sequence depth scaling [25]. This normalization step removed differences in sequencing depth between replicates by scaling each sample to an average depth of 1x.

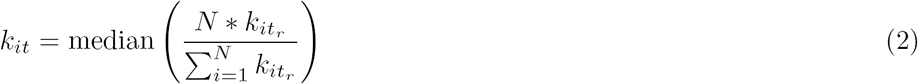

After aggregation, the combined signals were normalized once more to scale any imbalances in replicate numbers back to 1x, prior to making comparisons between replication times.

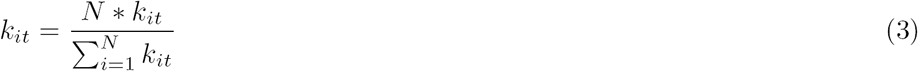

Our *A. thaliana* test data was relatively high coverage at 30x per bioreplicate, so we used the median function to generate a robust signal, instead of defaulting to sum.

### Reducing Type I Error

Repliscan aims to detect and highlight peaks of replication coverage, but some peaks may be too high and may in fact be false-positives caused by errors in the reference. For instance, if a repetitive element is present three times in the actual genome, but present only once in the reference sequence due to assembly error, all reads would align uniquely to the same location. If two of the actual elements replicate early and the third in middle S phase, the early peak would be twice as large and dominate the classification process. To reduce type I error arising from genomic repeats, we needed to detect and exclude these areas from the final classification because there is no way to resolve such duplication events without improving the reference genome.

The distribution of sequencing coverage is bounded on the left at zero, with very long, positive tails (Figure 4). Before we can detect any outliers we first need to transform the data to actually fit a probability distribution.

In Figure 4, we show that both the log and square root transformations stabilized the spread and skew. The log transformation extends the (0,1) tail and shortens the [1,∞) tail, making the distribution more normal-like. The square root transform also shortens [1,∞) tail and spreads the [0,1) tail, but not to the same extent, leaving the distribution skewed towards 0. While different, both transformations improve the fit of different probability distributions.

**Figure 4.**
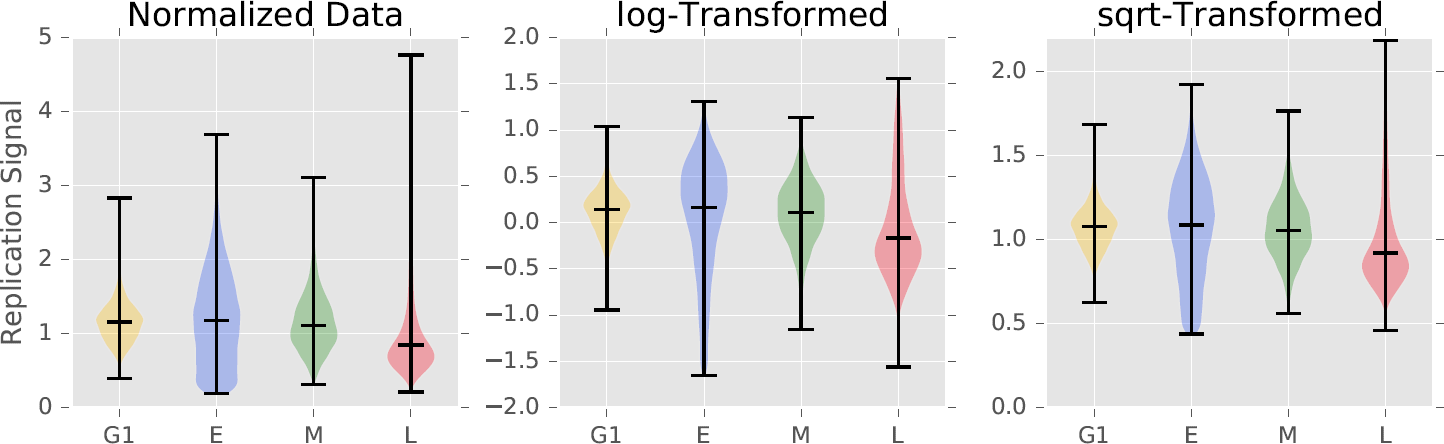
Normalized and transformed replication signals. Violin plots showing how the normalized and aggregated *A. thaliana* chromosome 3 replication signals from G1, early (E), middle (M), and late (L) S-phase data was bounded from [0,1). We separately experimented with with log transforms to make the distributions more normal-like, and square root transforms to stabilize the spread.

Normally, sequencing depth is modeled with a Poisson distribution because the integer counts are discrete[26], positive, and asymmetric. However, our aggregated and normalized data is continuous, positive, and asymmetric. To accurately model these sequencing values we use the Gamma distribution for highly-skewed data and the normal-like methods for symmetric data[27]. In all, we provide four combinations of methods to transform the data and detect outliers:

fitting a gamma distribution to the log transformed data,

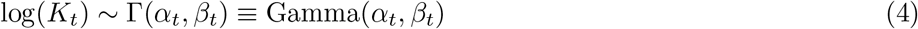

fitting a gamma distribution to the square root transformed data,

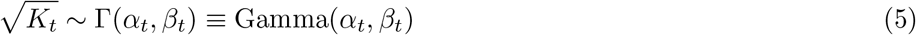

fitting a normal distribution to the log transformed data,

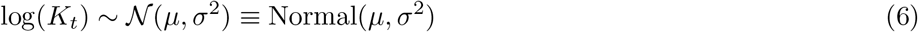

or calculating the whisker bounds (WB) of a boxplot from the log transformed data

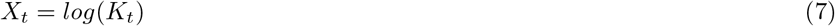

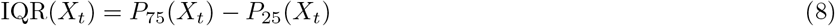

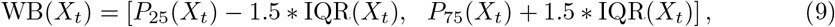

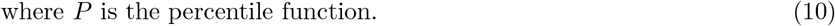

We use scipy[28] version 0.15.0 to fit all probability distributions to the actual coverage windows. Windows with coverage in the upper and lower 2.5% tails of the calculated probability distributions, or outliers when using whiskers, are considered unrepresentative and removed (Figure 5).

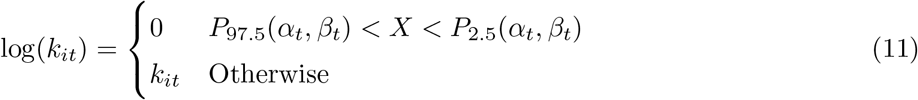

For simple cases, or when the transformed data does not resemble a probability distribution, we also provide the option of a rank-based (percentile) cutoff. By default, this will remove the upper and lower 2.5% coverage values, but this value can also be customized by the user.

The outliers in the positive coverage tails that this method removes may comprise a significant amount of coverage, so we perform another round of normalization to return the sample to 1x coverage. Each of the five methods has its own strengths and computation complexity. Most coverage data can be accurately modeled with the normal distribution. For cases when the transformed coverage distributions are still skewed, we suggest using the gamma distributions. If for some reason, the coverage data is multimodal, the whisker or percentile cutoff methods will both remove outliers from the data. We recommend the whisker method over a percentile cutoff because the whiskers remove data from a derived distribution, while the percentile indiscriminately removes a percentage of the data.

**Figure 5.**
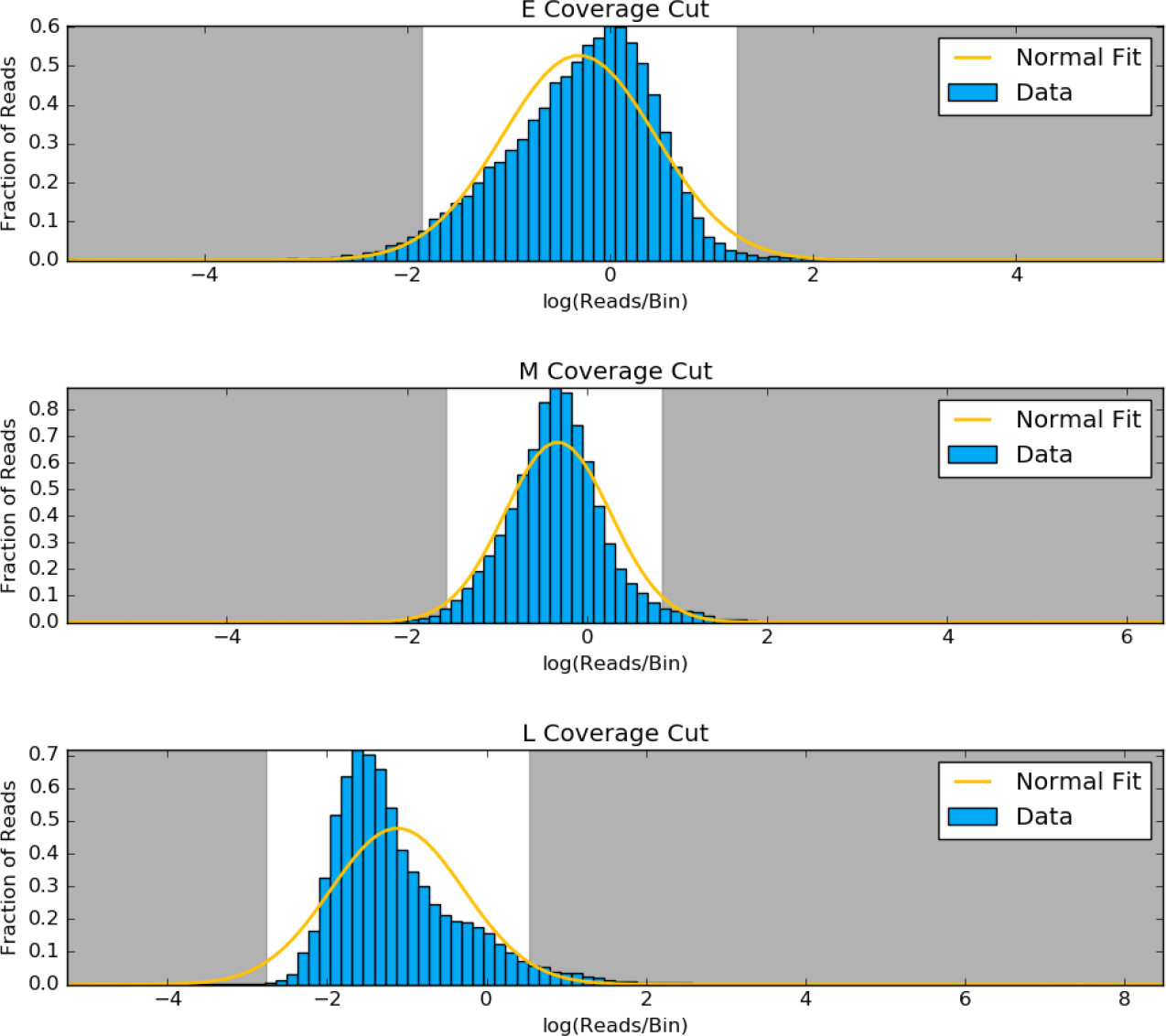
Outlying coverage in chromosome 3. Based on the normal distribution fit (yellow) to the log transformed coverage distribution of early (E), middle (M), and late (L) S-phase data, windows that fall in the tails shaded in gray are removed from the analysis.

### Normalize for Sequenceability

Amplification, fragmentation, and shotgun sequencing DNA is a non-uniform random process. Coupled with imperfect alignment efficiency from repetitive regions and incomplete reference genomes, artificial peaks arising from differences in the efficiency with which specific genomic regions can be sequenced are easy to confuse with actual signal peaks. This does not have a significant impact on comparisons between samples, but makes it difficult to compare adjacent genomic regions. Our sequencing protocol included a sample of non-replicating G1 DNA to correct for this phenomenon.

In G1, the cell is growing in physical size but no DNA replication is taking place, so the copy number of each sequence in the genome is at the 2C level. Variations in sequenceability can thus be separated from variations in signal attributable to DNA replication. Dividing each of the S-phase samples by the G1 sample normalizes each of the windows by giving the ratio of treatment coverage over expected coverage.

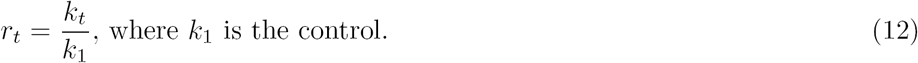

To better illustrate this process, consider two replication coverage windows next to each other: the first one is accessible and easy to sequence, and therefore produces more fragments per unit input DNA than the second window, which is hard to sequence. The normalization step would lower the signal from the first window, dividing it by a big coverage number from G1. It would also raise the signal from the second window, which would be divided by a smaller G1 number, making the two windows more comparable and reducing background noise. We recommend that such a control be implemented in all DNA sequencing based experiments to detect replication timing, on the basis that a non-replicating G1 control is the best, and most uniform representation of the genome. However, in the event that a non-replicating G1 is not sequenced, all S-phase samples can be combined to synthesize a total-S control, or a total DNA control can be used.

### Haar Wavelet Smoothing

Data sampling is always affected by noise. Statistical noise can be accounted for and modeled with more sampling, more robust statistical methods, or by summarizing larger ranges of data. Adding replicates for additional statistical power is cost-prohibitive, especially for larger genomes. Instead, we adopted the Haar wavelet transform to summarize replication data as an orthonormal series generated by the Haar wavelet. Using Wavelets[29] version 1.0, we performed a maximum overlap discrete wavelet transform with the Haar wavelet using reflected boundaries and level 3 smoothing on a per-chromosome basis for each sample. Wavelet decomposition is designed to represent a signal as a collection of frequencies. Level 3 decomposition represents a signal as the upper 87.5% of frequencies. Smoothing works as a low-pass filter, where small and frequent changes are removed, while large and wide changes are preserved.

We specifically chose the Haar wavelet over other smoothing methods because it is a square function with discrete boundaries and thus resembles the signals we aim to detect. General smoothing methods like LOESS and moving average methods produce stabilized trends from data, but they work by summarizing subsets of the whole picture. These methods also leave behind artifacts. A moving average will change a square peak into a sawtooth pattern the size of the smoothing window and will be affected by a single point of noise. LOESS is designed to model trends in sliding subsets of the data, but each of the least-squares regression steps are vulnerable to noise as with the moving average. LOESS will also spread out peaks in our data because of our uniform window size (1 kilobase), and is designed to accurately model clusters of points. As demonstrated in Figure 6A with simulated data, the Haar wavelet accurately removes low-amplitude and high-frequency noise to reconstruct the original signal without artificially expanding the peaks of replication signal. Applying the moving average, LOESS, and Haar wavelet to actual *A. thaliana* data in Figure 6B shows that both the moving average and LOESS can capture large trends, but the Haar wavelet excels at highlighting subtle peaks in the data without under smoothing and requiring the user to choose the range they summarize on. Any proportion or range of the data is very different when choosing different window sizes. Haar only removes low-amplitude frequency trends from the wavelet transform.

**Figure 6.**
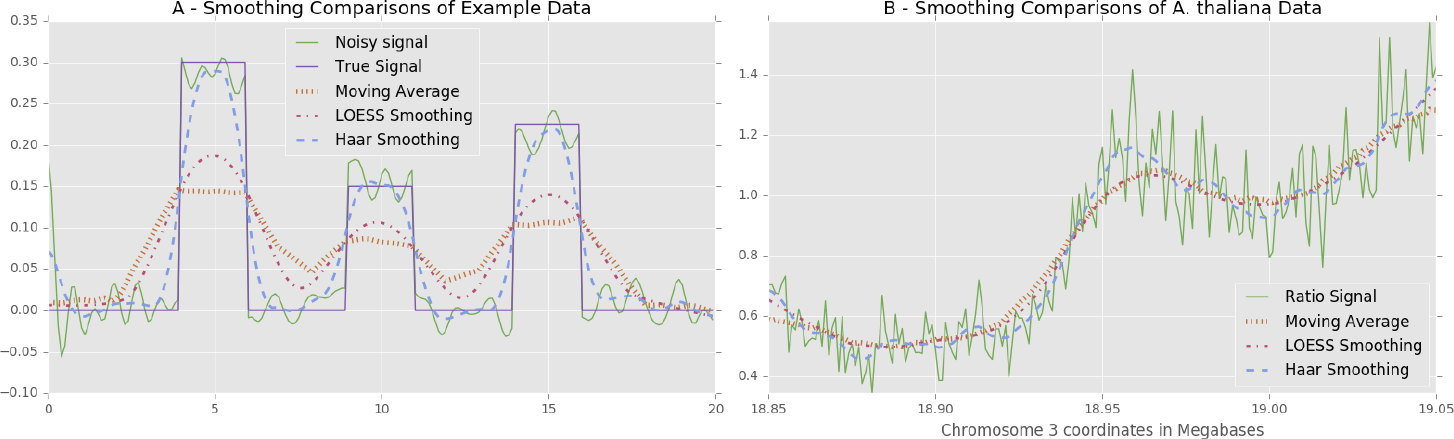
Smoothing comparisons. **A** – Noise (green) is added to an original signal (purple), and then smoothed with a 4 unit (40 point) moving average (orange), a 5 unit (25% subset) LOESS (red), and a level 3 Haar wavelet (blue). Both the moving average and LOESS spread out the peaks and artificially lowered signal amplitudes, while the Haar wavelet keeps bounds and peak heights close to the original. B – The *A. thaliana* middle S-phase normalized signal (green), is smoothed with a moving average (orange), LOESS (red), and the level 3 Haar wavelet (blue) for comparison.

We experimented with several levels of decomposition with our data, and found that the low-frequency trends preserved with level 3 aligned to genes, transposable elements, and histone marks on each genome the best. If the window size is kept at the default of 1 kilobase, this decomposition level can be kept the same because the same frequencies are represented. If the window size is changed to accommodate different sequencing depths, we suggest that users experiment with different decomposition levels, because this essentially changes the sampling rate of the analysis.

### Defining Replication

The analysis to this point yields a smoothed ratio of normalized replication ratio signals *r_cwt_* in windows (*w* = 1..*Y*) per chromosome (*c* = 1..*X*), with a range of [0,∞) that can be compared to each other, and leads to the question of which signals can be considered confidently as resulting from DNA replication. Lee *et al.* [15] originally considered array-based replication signals greater than the control as actively replicating in their investigation of *A. thaliana* as follows.

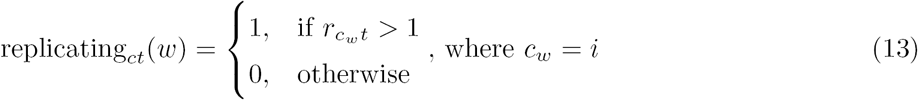

The Repliscan software allows users to adopt this threshold method, but we also include more robust methods to define replication. The simple threshold approach above is appropriate when considering replication as a ratio, but because all signals from the early, middle, and late S-phase samples represent labeled – and therefore, replicating – DNA, even signals that are less than the control must be considered as reflecting some level of replication activity. In other words, even though there may be noise in the data, all replication signals should be genuine because EdU is only incorporated into newly replicated DNA. Instead of simply choosing a smaller ratio threshold, we implemented a percentile cutoff based on the distribution of the ratios. By default, this method removes the lowest 2% of the values for a chromosome in a given sample.

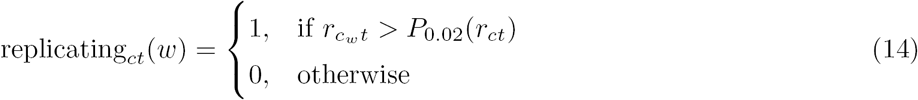

While this method is a data-dependent means for establishing a cutoff, it was not considered ideal for an automatic analysis for two reasons. First, a cutoff is still being dictated, even if it is more robustly supported than in previous analyses. Second, this cutoff will always remove a flat percentage of the values, even if there is no evidence they are not high-quality data points. To improve on these deficiencies, we implemented a threshold for replication that depends on the information provided in addition to the data.

To maximize the fraction of a chromosome with valid replication signal (or information), we designed an optimization method that incorporates as much of each chromosome as possible by analyzing the rate that chromosome coverage changes with replication signal. Using data from all time points, coverage is defined as the fraction of windows with a signal greater than the threshold in at least one replication time.

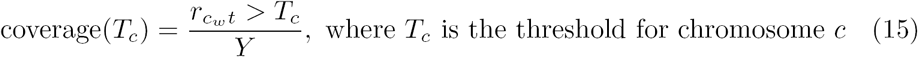

Our optimization process begins from the point of the largest absolute change in coverage (*mT_c_*), and lowers the replication threshold (*T_c_*) until the absolute chromosome fraction per sample/control coverage differential goes below 0.1, effectively leveling out. In rare cases where this process does not converge, the threshold is set to be the median of all chromosomes that do converge.

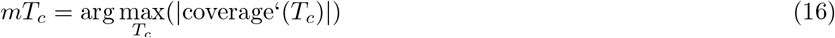

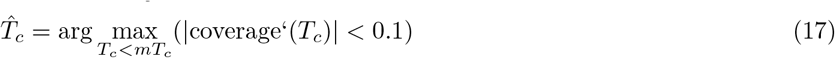

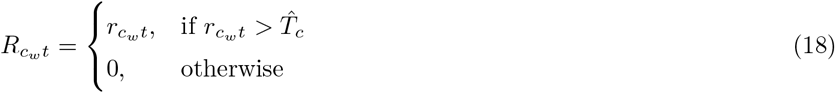

Such a search pattern circumvents any local optima in the coverage signal that may have stalled a gradient descent. That being said, we implemented the threshold to run on a per-chromosome basis to minimize the effect of any structural differences (Figure 7).

**Figure 7.**
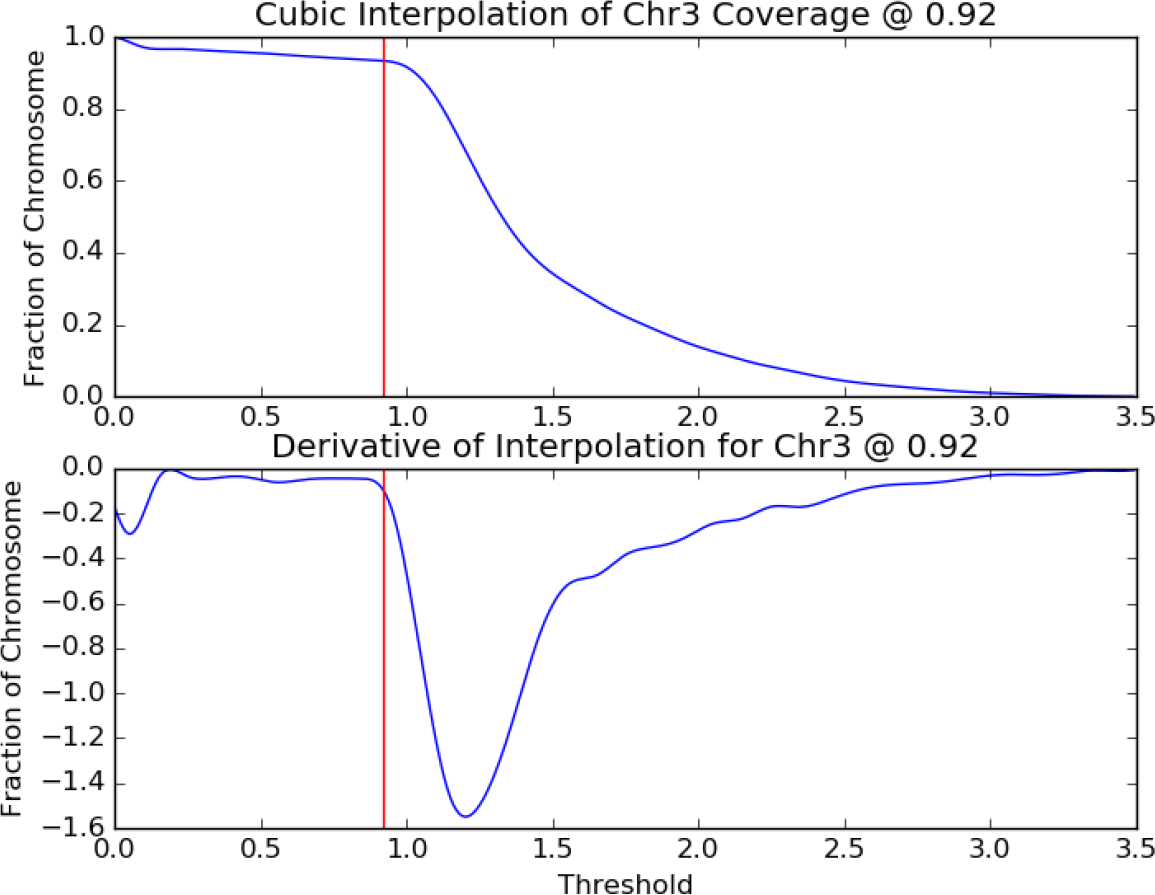
Replication threshold from coverage. The upper plot shows how much of *A. thaliana* chromosome 3 will be kept for downstream analysis as a function of the signal threshold. The lower plot shows the chromosome coverage differential as a function of the threshold. The vertical red line in each plot marks the optimal threshold of 0.92.

The end result is a method that includes as much of the genome and coverage information as possible, and prevents the use of small signals when they comprise a small portion of the chromosomes. Our method is generically applicable to experiments using the same Repli-seq protocol because the threshold is calculated from the data. A critical benefit is that users are not required to be masters of their data or this tool, and can instead focus on interpretation.

### Classification/Segmentation

Given a signal that can confidently be considered as arising from DNA replication, we are able to classify segments of the genome according to when in the cell cycle they are replicated. Suppose that in one of the windows in Chromosome 3, we have the following levels of replication in Table 1.

**Table 1.** Example coverage values to demonstrate replication timing classification.

| Time | Early | Middle | Late |
| --- | --- | --- | --- |
| Coverage | 0.93 | 0.8 | 3.0 |
| Replicating | 0.93 | 0 | 3.0 |

We already know from Figure 7 that any values below 0.92 in Chromosome 3 are not considered replicating, so the middle S-phase value would become 0 and we would say this window replicates in both early and late S-phase. However, the late replication level is 3 times higher than that of early, which is just past the threshold for replication at 0.93. Instead of making another replication threshold, we implemented a general solution to compare values against each other using a proportion.

First, on a window-by-window basis, we take the infinity norm of all values, which means we divide all values by the maximum for that window position.

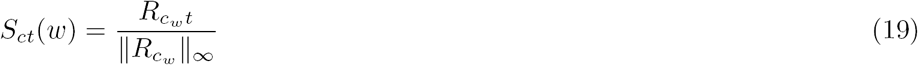

This operation scales the largest value to 1 and the others to the range [0,1]. A time signal is then classified as predominantly replicating *C_c_t*(*w*) if the normalized value is greater than 0.5, or at least half the size of the largest signal for that window.

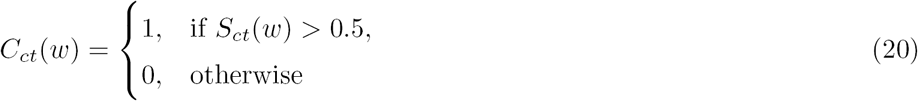

The infinity-norm ensures that the largest value will always be classified as replicating, and this classification method allows for a window to be called strongly replicating at more than one time in S-phase (e.g. both early and late) when other signals are within 50% of the maximum value. Besides 0.5 being easy to test for, this creates an equally partitioned solution space in the form of an n-dimensional hypercube. In the case of our *A. thaliana* data, the space is a 3-dimensional cube with each dimension being one of the time points: early, middle, and late S-phase. The 0.5 partition then creates 8 equal-sized sub-cubes corresponding to each possible combination of times:

{Non-replicating, Early, Middle, and Late}

along with

{Early-Middle, Middle-Late, Early-Late, and Early-Middle-Late}

S-phase replication combinations.

## Results and Discussion

### Data

To demonstrate the ability of our methods to adapt to different datasets, we ran our pipeline on the *A. thaliana* Col-0 cell culture data (PRJNA330547) that was used to develop these methods, and a separate similarly prepared *Z. mays* B73 replication timing dataset (PRJNA327875) also from our lab.

#### A.thaliana

The *A. thaliana* experiment was comprised of 3 early S bioreplicates, 3 middle S bioreplicates, 3 late S bioreplicates, and 1 G1 sample. Each bioreplicate was paired-end sequenced to 36x coverage. The unique and properly-paired alignment rate for each sample was approximately 85%, yielding a total of 30x viable replication data from each sample. Due to the high coverage, we decided to use 1 kilobase windows and merge bioreplicates with the median function for our analysis.

#### Z. mays

In the *Z. mays* experiment, there were 3 early S bioreplicates, 3 middle S bioreplicates, 2 late S bioreplicates, and 2 G1 technical replicates. Each bioreplicate was paired-end sequenced to about 5x coverage. While there were more reads than the *A. thaliana* experiment, the *Z. mays* genome is much larger, so coverage was lower. Using the B73 AGPv3 genome assembly, the unique and properly-paired alignment rate for each sample was approximately 99%, yielding a total of 5x viable replication data from each bioreplicate. Even though a larger analysis window could have been used, we decided to use the same 1 kilobase windows for this dataset, and deemed the summation of bioreplicates was necessary to achieve enough coverage to highlight peaks in the data.

### Segmentation Overview

Using 1 kilobase windows, median aggregation for *A. thaliana*, and sum aggregation for *Z. mays*, we used our default pipeline to classify the replication timing of our data. We generated Figure 8 to show the replication segmentation classification of Chromosome 3 in *A. thaliana* and Chromosome 10 in *Z. mays*.

**Figure 8.**
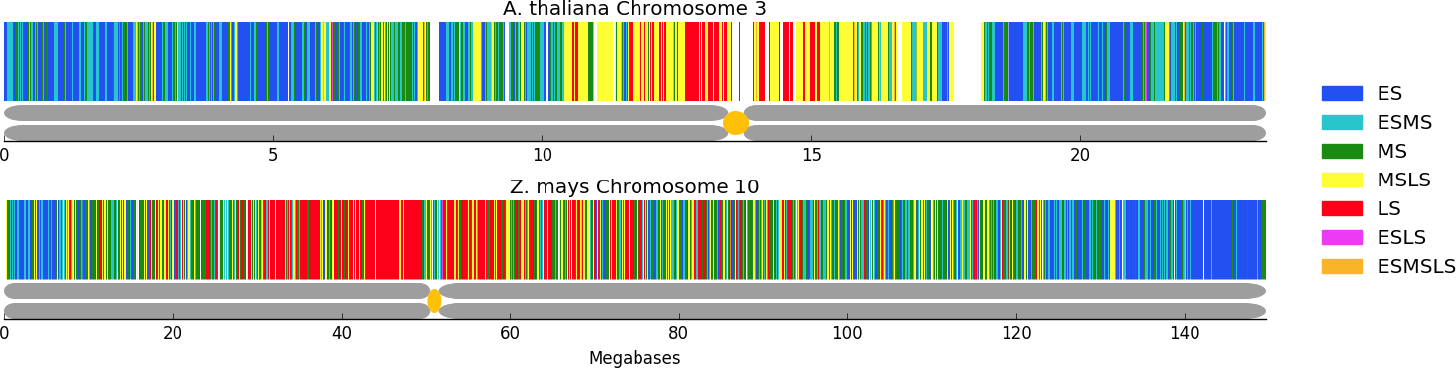
Comparison of A. thaliana and Z. mays segmentation. Following the segmentation legend on the right, *A. thaliana* chromosome 3 (top) and *Z. mays* chromosome 10 (bottom) have been classified into segmentation regions by Repliscan. The large white regions in the *A. thaliana* figure are unclassified regions due to high or very low signal. Below each replication segmentation is a depiction of the chromosome, with the centromere location marked in yellow [30, 31].

In both instances, early replication is concentrated toward the ends of the chromosome arms, with middle and late replication becoming more prominent closer to the centromere and the highest concentration of late replicating sequences in the heterochromatin surrounding the centromere. These timing maps demonstrate that the method developed using the *A. thaliana* data was successfully applied to the lower coverage *Z. mays* data by simply choosing to aggregate replicates using the sum instead of the median.

### Segment Composition and Size

Instead of viewing the chromosomes as a whole, we can also get an idea of predominant replication times by looking at the proportional composition. Figure 9 shows that Early, Early-Middle, and Middle-Late S-phase replication makes up most of the segmentation profiles for *A. thaliana* Chromosome 3. About 6% of the chromosome is missing around the centromere and heterochromatic knob, which probably would have been classified in the Middle to Late times based on what we do see. In *Z. mays*, we see a more uniform distribution of Chromosome 10, which is 5-fold larger, across the replication segmentation classes. Lee *et al.* [15] previously hypothesised a two-stage replication program, but our results, which were generated using much shorter labeling times to capture much smaller increments of replication, show a more even spread (Figure 9).

**Figure 9.**
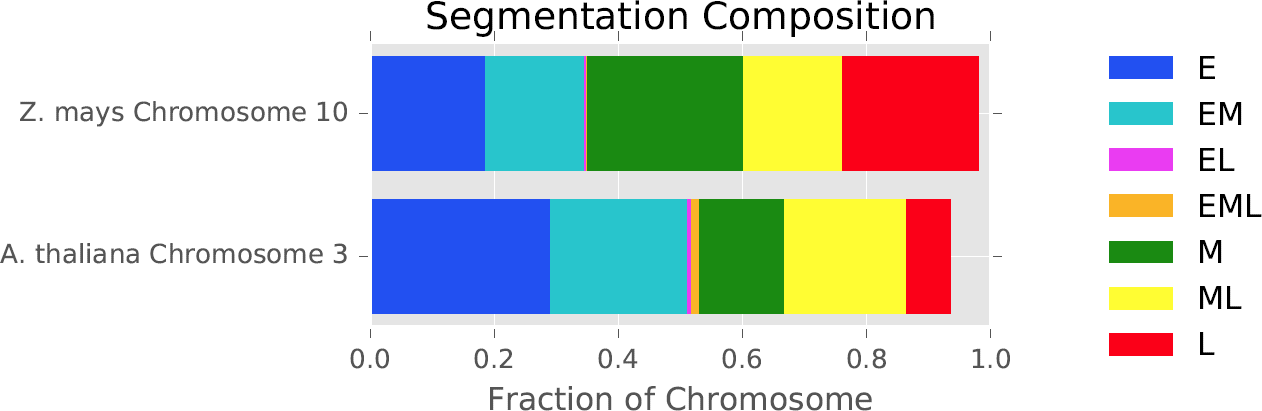
Composition of replication segmentation. The segment composition shows that replication in *A. thaliana* is skewed towards early S replication, while *Z. mays* has an even distribution across early, middle, and late S. We can also see that the non-sequential early-late (EL) and early-middle-late (EML) classifications comprise a very small proportion of the classified segments in both cases.

The Early-Late and Early-Middle-Late comprise a small portion of the chromosomes in both organisms and could arise naturally in the data through allelic and cell population differences. Figure 10 shows a different summary of the segmentation breakdown, highlighting the segment size distribution with boxplots. Once again, Early-Late and Early-Middle-Late segments are distinct in that their lengths are small relative to the other timing categories.

### Downsampling and Stability of Results

The relatively small genome size of *A. thaliana* allowed us to obtain extremely deep sequencing coverage, which is currently cost-prohibitive for larger genomes. To estimate a minimum coverage requirement for our methods, we simulated experiments with lower coverage via downsampling. We first generated 3 technical replicates by randomly sorting the original alignment files. We removed reads from each of the replicates in 1% increments without replacement. Each of the 300 (100 x 3) simulated experiments were analyzed using both median and sum aggregation, and no (none), log gamma, square root gamma, normal, and whisker outlier removal. To account for differences arising from the sorting order, the final classification for each window was determined by majority across the 3 replicates. Classification ties were broken by treating the early, middle, and late time classification combination as a 2-bit binary number, and taking the median.

**Figure 10.**
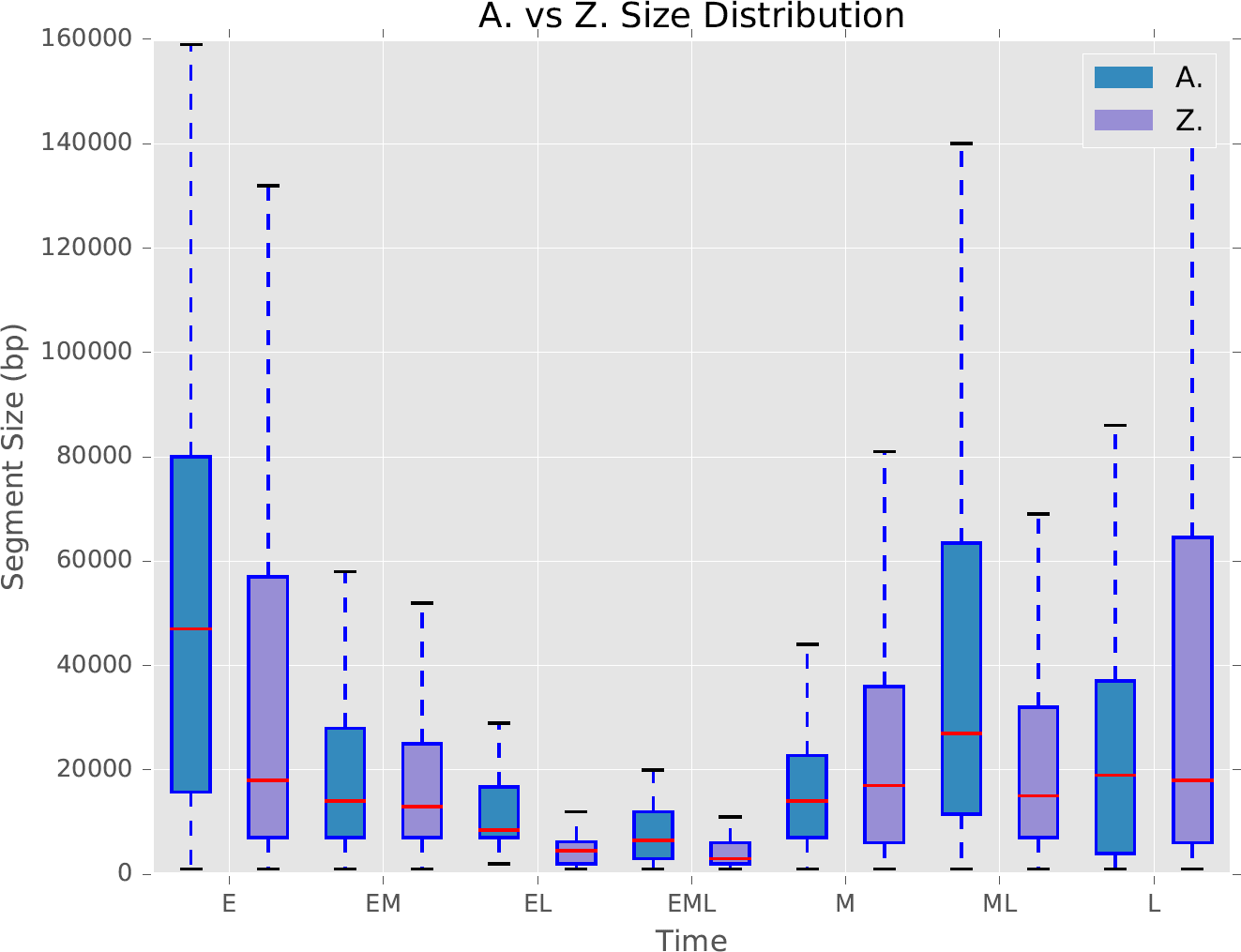
Segment size distribution. Boxplots for every combination of replication time, illustrating the distribution of segment sizes. Early (E) and mid-late (ML) S were largest in *A. thaliana*, while early and late (L) were largest in *Z. mays*.

**Figure 11.**
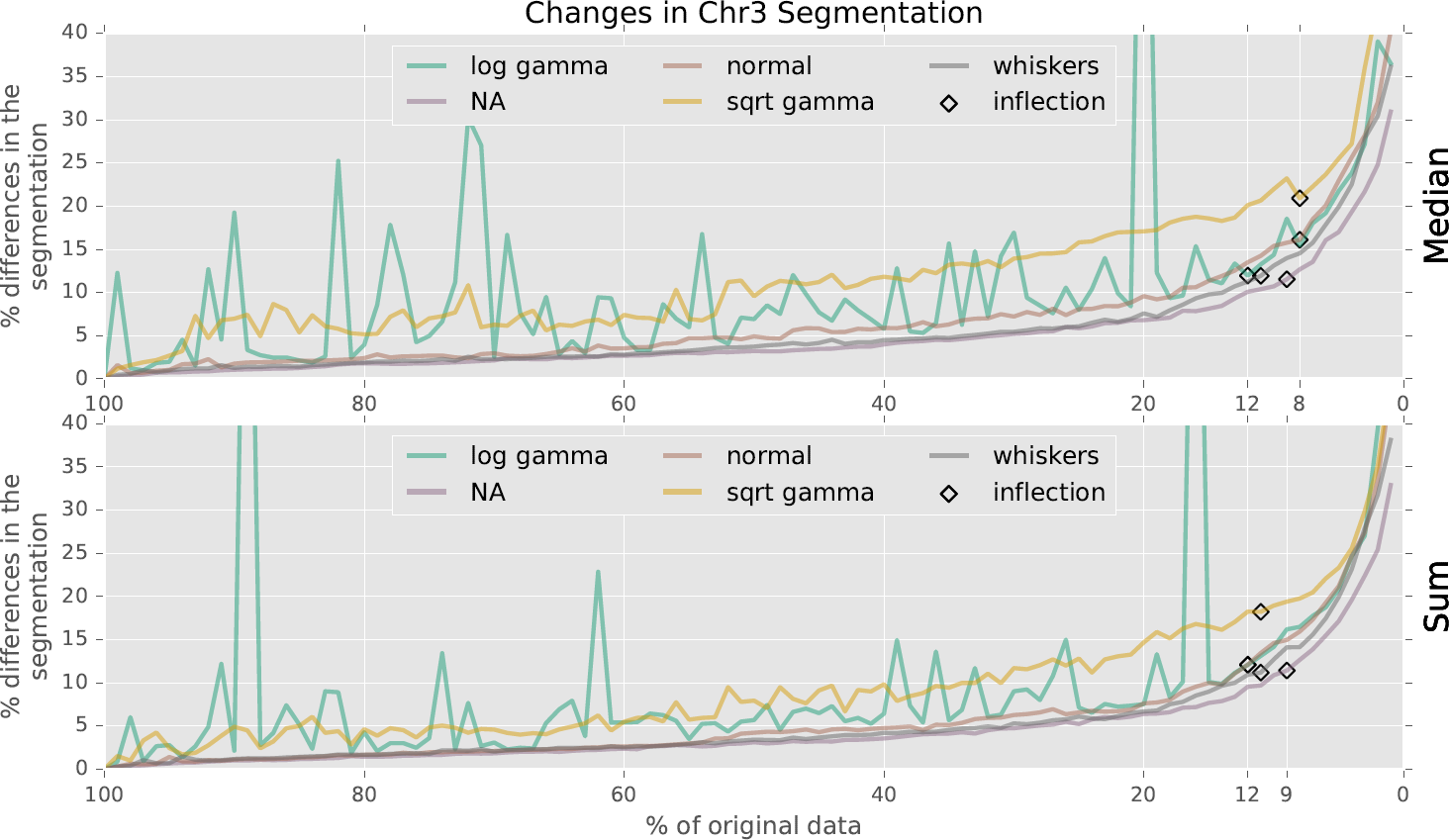
Segmentation differences in downsampled data. After downsampling the *A. thaliana* data, the accuracy of median (top) and sum (bottom) aggregation, and outlier detection using log gamma, none (NA), normal, square root gamma, and whiskers. Inflection points in the differences are labeled with black diamonds.

After confirming that the segmentation profiles from all three 100% replicates were identical to our original segmentation, differences for each run type were calculated as percent Hamming distances from the 100% version. All differences were compounded and plotted as a fraction of the whole chromosome in Figure 11. The most obvious results are the spikes of differences in both the median and sum log transformed gamma runs when the iterative fitting function failed to converge (Figure 12).

**Figure 12.**
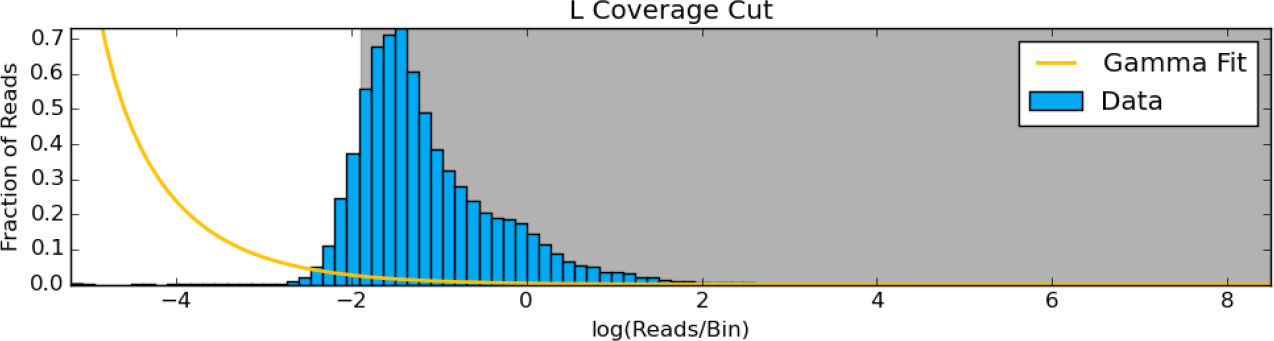
Unconverged log gamma fit. Most of the data is removed when the iterative fitting function fails to converge with the log transformed gamma distribution. Instances like this produce the spikes of differences in Figure 11.

Shifting attention to the square root gamma experiments in Figure 11, we see that the fit function never fails to converge, but there is increased variability of results among each level of downsampling. All other probability functions are very stable between downsampling runs. We even see that summing the coverage to 90x provides no improvement over the median – even at low coverage levels. The inflection points show that the most stable method was aggregating replicates with the median operation and removing coverage by fitting a normal distribution to the log transformed data. Results from this method began to noticeably diverge when downsampled to 8%, or 2.4x coverage. This indicates 5x coverage for the commonly studied species *Z. mays* (2.3 gigabase genome[32]) is sufficient to calculate a replication profile, which is quite tractable for a laboratory of modest financial means.

### General Application of Repliscan

To demonstrate that Repliscan is generally applicable, we used it to analyze two published Repli-seq datasets: Human fibroblast data from Hansen *et al.* 2010[12] (GSM923444) and *D. melanogaster* data from Lubelsky *et al.* 2014[33] (PRJNA63463).

The Human fibroblast Repli-seq data contains samples from 6 fractions of S phase (G1b, S1, S2, S3, S4, and G2) with two replicates each providing an average depth of 0.02x coverage. Using the supplementary methods of Hansen *et al.*, we were able to reproduce their original tag density results. Reads from both replicates were first combined and then aligned to the human reference genome (hg19). After alignment, signals with more than 4 reads per 150 basepair window were removed. Lastly, a percent total coverage in 50 kilobase wide windows was calculated every 1 kilobase (Figure 13).

To analyze this data with Repliscan starting from the aligned reads, we first needed a sequencing control. Both G1b and G2 contain replicating DNA in this experiment, so we combined G1b, S1-4, and G2 to create a total-S (TS) control in the first line of the Repliscan input configuration. After crafting the configuration file, we ran Repliscan with a window size of 50 kilobases and aggregation through sum to match the methods of Hansen *et al.* Figure 13 compares the output of Repliscan against the reproduced results in a region from their original work. Given that there were 6 fractions of S-phase in the Repliscan input, there were (2^6^ *−* 1) 63 possible classifications, but only 22 were present in the output segmentation. Repliscan presented temporally sensible results with replication initiating in G1b and spreading to G2 all while relying on the automatic tuning of Repliscan after matching the window size (Figure 13). We compared our results from Repliscan to the “BJ-G1 segment” regions published by Hansen *et al.* in their Supplementary Table S4 using the accuracy statistical measure.

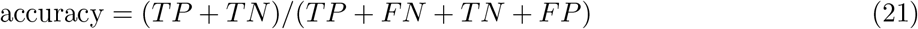

Where *T P* is the number of G1bS1 Repliscan classifications that match “BJG1 segment”, *F N* is the number of non-G1bS1 classifications that match “BJG1 segment”, *T N* is the number of non-G1bS1 classifications that also do not match “BJ-G1 segment”, and *F P* is the number of G1bS1 classifications that do not match “BJ-G1 segment.” We found that our Repliscan reanalysis had an accuracy of 83% with the published “BJ-G1 segment” results.

**Figure 13.**
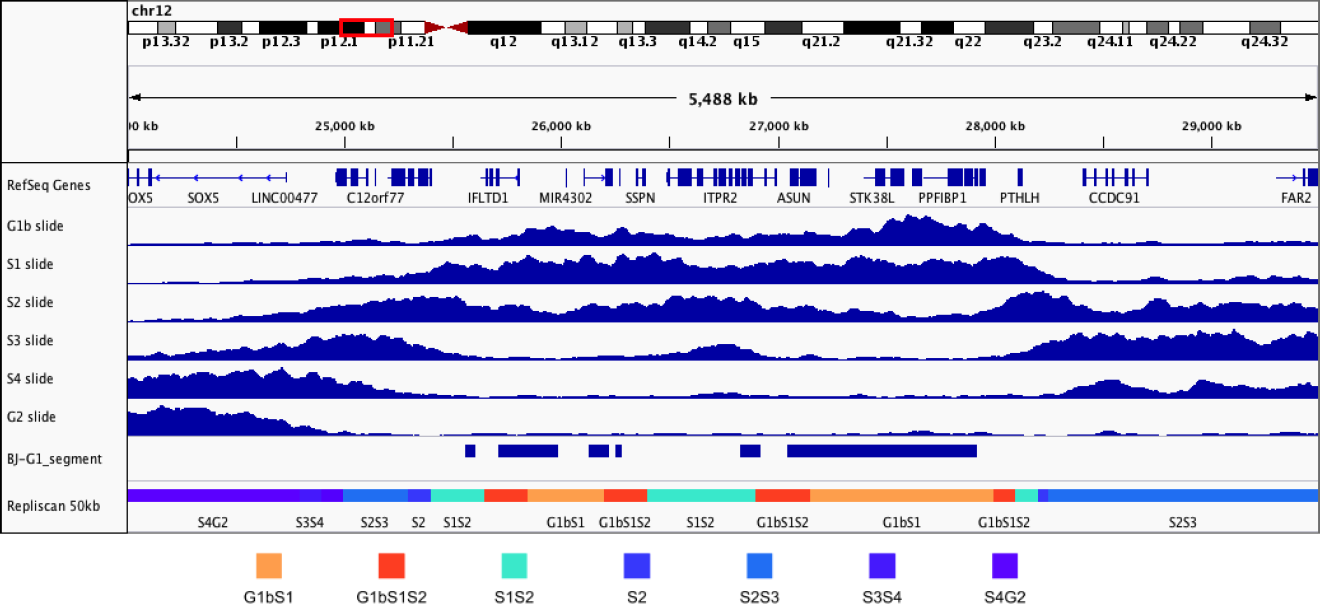
Human fibroblast Repli-seq. 50 kilobase sliding window replication signals (blue) reproduced from Hansen *et al.*, published “BJ-G1 segment” regions, and 50 kilobase Repliscan results (bottom).

We also reproduced the original continuous replication profiles of Lubelsky *et al.* by processing the raw data as was done in the original paper. Replicates were combined from each fraction of S phase (Early, Early-Mid, Late-Mid, and Late) and aligned to the dm3 Release 5.12 genome. Unique alignments were kept and the RPKM was calculated in 10 kilobase windows along the genome. The RPKMs from the 4 samples were then weighted and combined to create a single replication signal from 0 to 1. The replication signal was then LOESS smoothed with a span of 200 kilobases (20 bins). This continuous signal was then classified as early replication when the value was less than or equal to 0.5, and late replication when above 0.5 (Figure 14).

Similar to the work by Hansen *et al.*, this experiment did not contain an non-replicating G1 control, so we combined all fractions into a total-S (TS) control. For inputting the raw data into Repliscan, we crafted two input configurations: one with Early (early, early-mid) and Late (mid-late, late) (2S) to match the discrete results of Lubelsky *et al.*, and another with Early, Early-Mid, Mid-Late, and Late classifications (4S) to highlight the classification capabilities of Repliscan. Coverage averaged around 4.4x, so we ran Repliscan with both (2S and 4S) input configurations, sum replicate aggregation, and 10 kilobase windows to match the original analysis (Figure 14).

**Figure 14.**
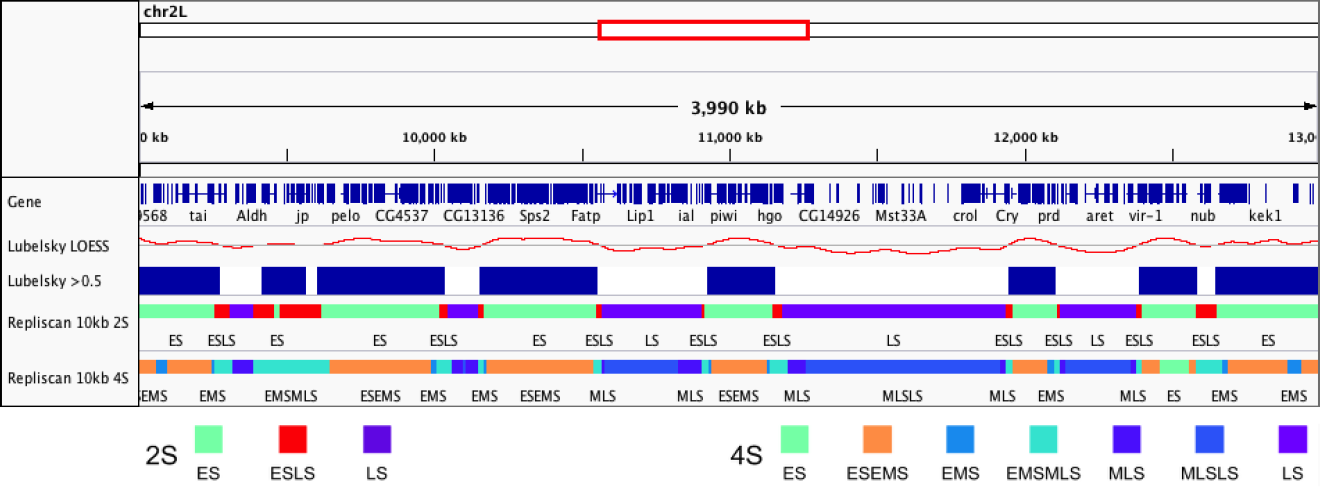
D. melanogaster KC167 Repli-Seq. Reproduction of the LOESS smoothed continuous replication profile (Lubelsky LOESS), and the thresholded, discrete early (blue) and late timing domains (Lubelsky *>* 0.5) from original Lubelsky *et al.* study. Repliscan segmentation results with Early (Early, Early-Mid) and Late (Mid-Late, Late) replication (2S), and Early, Early-Mid, Mid-Late, and Late replication (4S) configuration with 10 kilobase windows.

The Repliscan configuration with two S-phase fractions (2S) highly resembled the thresholded continuous signal (Lubelsky *>* 0.5) with a statistical accuracy measure of 95%. When Repliscan was run to capture all 4 S-phase combinations, more information was revealed about the replication timeline. Looking at the two left-most late regions of “Lubelsky *>* 0.5” in Figure 14, we can see that the continuous signal rides along the 0.5 threshold, and Repliscan predicted a long region of EMS-EMLS with all four fractions of S taken into context, instead of detecting an initiation site in the center. This situation is a good example of the type of coarse grained calls that we are trying to avoid with Repliscan by allowing combinations of replication in our classifications. Our 4S results were also found to be highly similar with the discrete data, with a statistical accuracy of 78%.

## Conclusions

Based on our results from running Repliscan on both *A. thaliana* and *Z. mays* data, we have demonstrated that our methods offer a robust means of analyzing data from replication timing experiments that use label incorporation. Although we argue that a non-replicating G1 control should be preferred for biological reasons, our analytical method can be used equally well with control datasets derived from synthetic total S phase pools or from total DNA. We have significantly improved on previous methods by incorporating non-destructive Haar smoothing, using optimization methods to define replication, and classification through signal proportion. When run using the same parameters but using data from different organisms, the methods automatically tuned their thresholds to adjust for differences in coverage. Downsampling our data showed our methods provided stable results at as little as 2.4x coverage and 1 kilobase analysis windows. Even lower coverages can be accommodated at lower resolution by using larger window sizes for the analysis. We also demonstrated that Repliscan can be used to classify replication regions in external Repli-seq data by applying it to both low-coverage Human and high-coverage *D. Melanogaster* experiments with 4 to 6 S-phase fractions and synthetic total-S controls. There is no current consensus pipeline for validation, so we compared the published results from the external datasets to those from Repliscan. We found that the Repliscan results were on average 85% identical to the original findings of these papers.

In-depth explorations of the replication programs in *A. thaliana* and maize will be published separately. We think these methods provide a path for greater understanding of the DNA replication program in plants, humans, and other higher organisms.

## List of Abbreviations

**TimEx:** Time of replication

**Repli-seq:** Replication label incorporation sequencing

**Edu:** 5-Ethynyl-2’-deoxyuridine

**BrdU:** 5-Bromo-2’-deoxyuridine

**NGS:** Next generation sequencing

**SRA:** Sequence read archive

**G1:** Gap 1 of cell division

**G2:** Gap 2 of cell division

**S:** Synthesis phase of cell division

**E:** Early S-phase replication

**M:** Middle S-phase replication

**L:** Late S-phase replication

**WB:** Whisker bounds

## Availability and requirements

Project name: Repliscan

Project home page: https://github.com/zyndagj/repliscan

Operating systems: Linux, OS X

Programming languages: Python v2.7

Other requirements: scipy v0.15.0+, samtools, bedtools v2.24.0+, wavelets v1.0, numpy, matplotlib.

## Declarations

### Ethics approval and consent to participate

Not applicable

### Consent for publication

Not applicable

## Acknowledgements

Special thanks to everyone in the Hanley-Bowdoin and Thompson labs.

## Availability of data and materials

The datasets supporting the conclusions of this article are available in the NCBI Sequence Read Archive (SRA) BioProjects PRJNA330547, PRJNA327875, and PRJNA63463 and GEO dataset GSM923444. All reproduced Human and *D. Melanogaster* Repli-seq results can be generated and viewed as described in the Repliscan repository.

## Funding

This work is supported by NSF IOS-1025830 “Epigenome Dynamics During DNA Replication”.

## Competing interests

The authors declare that they have no competing interests.

## Author’s contributions

GJZ developed and implemented the algorithm, and wrote the manuscript. EEW, LC, WFT, and LHB provided biological expertise and produced the data. JS provided feedback on the methods and helped revise the manuscript. All authors read, helped revise, and approved the final manuscript.

